# Unsupervised data-driven stratification of mentalizing heterogeneity in autism

**DOI:** 10.1101/034454

**Authors:** Michael V. Lombardo, Meng-Chuan Lai, Bonnie Auyeung, Rosemary J. Holt, Carrie Allison, Paula Smith, Bhismadev Chakrabarti, Amber N. V. Ruigrok, John Suckling, Edward T. Bullmore, MRC AIMS Consortium, Christine Ecker, Michael C. Craig, Declan G. M. Murphy, Francesca Happé, Simon Baron-Cohen

**Affiliations:** Department of Psychology, University of Cyprus, Nicosia, Cyprus; Center for Applied Neuroscience, University of Cyprus, Nicosia, Cyprus; Autism Research Centre, Department of Psychiatry, University of Cambridge, Cambridge, UK; Child and Youth Mental Health Collaborative at the Centre for Addiction and Mental Health and The Hospital for Sick Children, Department of Psychiatry, University of Toronto, Toronto, Canada; Department of Psychiatry, National Taiwan University Hospital and College of Medicine, Taipei, Taiwan; School of Philosophy, Psychology and Language Sciences, Department of Psychology, University of Edinburgh, Edinburgh, UK; Centre for Integrative Neuroscience and Neurodynamics, School of Psychology & Clinical Language Sciences, University of Reading, Reading, UK; Brain Mapping Unit, Department of Psychiatry, University of Cambridge, Cambridge, UK; Cambridgeshire and Peterborough NHS Foundation Trust, Cambridge, United Kingdom; Sackler Institute for Translational Neurodevelopment, Department of Forensic and Neurodevelopmental Sciences, Institute of Psychiatry, Psychology and Neuroscience, King's College London, London, UK; Autism Research Group, Department of Psychiatry, University of Oxford; MRC Social, Genetic, and Developmental Psychiatry Centre, Institute of Psychiatry, Psychology & Neuroscience, King's College London, London, UK; Department of Child & Adolescent Psychiatry, Psychosomatics and Psychotherapy, University Hospital Frankfurt am Main, Goethe-University, Frankfurt am Main, Germany; National Autism Unit, Bethlem Royal Hospital, SLAM NHS Foundation Trust, UK

**Author notes:** Corresponding Author: Michael V. Lombardo, University of Cyprus, Department of Psychology, 1 Panepistimiou Avenue, Aglantzia, Nicosia 1678, Cyprus. Equal contributions (Michael V. Lombardo, Meng-Chuan Lai). Please see the full list of individuals within the MRC AIMS Consortium at the end of the manuscript.

**Keywords:** autism, mentalizing, emotion recognition, heterogeneity, precision medicine, clustering

## Abstract

Individuals affected by autism spectrum conditions (ASC) are considerably heterogeneous. Novel approaches are needed to parse this heterogeneity to enhance precision in clinical and translational research. Applying a clustering approach taken from genomics and systems biology on two large independent cognitive datasets of adults with and without ASC (n=715; n=251), we find replicable evidence for 5 discrete ASC subgroups that are highly differentiated in item-level performance on an explicit mentalizing task tapping ability to read complex emotion and mental states from the eye region of the face (Reading the Mind in the Eyes Test; RMET). Three subgroups comprising 42-65% of ASC adults show evidence for large impairments (Cohen’s d = −1.03 to −11.21), while other subgroups are effectively unimpaired. These findings delineate robust natural subdivisions within the ASC population that may allow for more individualized inferences and accelerate research towards precision medicine goals.

---

Autism spectrum conditions (ASC) are currently defined by consensus behavioral criteria of difficulties in social-communication and restricted repetitive behaviors. Although the population is subsumed under a single unitary diagnostic label, variability between affected individuals is considerable^1^. This diversity or ‘heterogeneity’ inherent within ASC can be seen at multiple levels, from a myriad of different etiological mechanisms^2^, developmental trajectories^3, 4^, sex/gender^5^, clinical comorbidities^6^, cognitive/behavioral features (e.g., language development^7^), and the list could go on^8^. Accordingly, many in the field now subscribe to the idea that there is not just one ‘autism’, but rather multiple ‘autisms’^9, 10^. Given these rich theoretical ideas about the complexity manifesting behind unitary clinical diagnostic labels like ‘autism’, it seems like a natural extension of logic that research would follow along such ideas. However, the opposite occurs a majority of the time, whereby clinical and translational research is done utilizing case-control comparison methodology that treats autism as one omnibus group and makes comparison to a matched ‘control’ or ‘comparison’ group. While the standard case-control approach appears to be congruent with the categorical divide expressed within psychiatric diagnostic manuals like DSM-5 and ICD-10, it is antithetical with the rich ideas suggesting that in order to move forward in understanding mechanisms affecting individuals with ASC, one cannot effectively utilize a paradigm that lumps together heterogeneous mixtures of different types of individuals. This problem is not necessarily specific to autism per se, and is a general issue within psychiatry that has prompted the rise of alternative approaches such as the NIMH Research Domain Criteria (RDoC)^11^ as well as ideas related to the concept of ‘stratified psychiatry’^12^. Closely related to these ideas are the more generalized goals of ‘precision medicine’ outlined for all domains of medicine^13, 14^ The goals of ‘precision medicine’ applied to ASC would outline an individualized approach to areas that can have immediate impact on treatment and support; from clinical assessment, diagnosis, personalized treatment and prognostic approaches, to better specificity in determining etiological mechanisms connected to specific phenotypes.

One primary challenge for meeting these goals is the basic question of how one should stratify a heterogeneous label like ‘autism’ into natural subdivisions that meaningfully point towards important underlying mechanisms and which could have potential for impact on clinical issues. There are multiple dimensions and levels through which one could start this process, from stratifications at the etiological level all the way up to subgroups distinguished by differing neural systems, cognitive, behavioral, and/or developmental patterns^7^. Analytically, stratification can be based on supervised knowledge driven by experimenter-based preconceptions, theory, and/or assumptions. In contrast to supervised approaches, data-driven unsupervised approaches can be advantageous in a variety of cases when there is limited a priori knowledge that can be used to supervise the stratification process. As for goals of the stratification process, we ultimately need stratification approaches that identify consistently replicable subgroups nested within the ASC population. Subgroup effect sizes should ideally naturally organize into much more robust patterns of clear difference or a lack thereof and such effects should also allow for much more parsimonious distinctions than standard case-control comparisons.

Stratification could be very important at the cognitive level, particularly when applied to cognitive phenomenon that links back to behavioral difficulties that are cardinal features of ASC^15^. Here we focus on the domain of mentalizing/theory of mind (henceforth ‘mentalizing’), which we have known for the last 30 years is a key cognitive explanation behind social-communicative difficulties in ASC^16, 17, 18^. Despite much progress over the last 3 decades, it is notable that a majority of the evidence to date rests on statistical evidence about what differs *on-average* in a case-control setting. Hidden within these on-average case-control differences is additional complexity at the individual level. Many individuals will show evidence of some kind of deficit in mentalizing over the lifespan, while others may not show any difficulty or may simply mask the difficulty via compensatory mechanisms^19, 20^. This heterogeneity will also likely change throughout development as individuals acquire more competence in the domain^21^. There are also conceptual distinctions within mentalizing, such as the distinction between explicit/controlled versus implicit/automatic processes, with the latter continuing to be atypical much later in life despite an individual possessing explicit abilities^20^. The concept of mentalizing has also expanded considerably over time from its more constrained initial usage that was much more closely tied to ‘theory of mind’ as measured by standard false belief tasks. The term is now notably quite broad with respect to the different kinds of social cognitive components that can contribute to the overall domain of mentalizing, such as processing eye gaze, recognizing emotion, intuitively tracking other’s mental status, inferring other’s mental status with efforts, separating belief from fact, comprehending social scripts, switching between self and other perspectives, etc. Thus, in addition to heterogeneity at the level of individuals, there is also conceptual heterogeneity what is considered ‘mentalizing’. Some have called for deconstruction of the domain of mentalizing into more basic components^22^ and we would agree that such steps are essential to enhancing the precision of our understanding on the topic.

In the current work, we focus precisely on components of mentalizing measured via a widely used task that involves explicit recognition of complex emotional and mental states simply from looking at the eye region of the face – the Reading the Mind in the Eyes Test (RMET)^23, 24^ In a recent study comparing a relatively large ASC sample to a typically-developing (TD) control group, we found on-average reduction in RMET performance. However, ASC distributions were notably negatively skewed and there was a significant degree of overlap of ASC individuals within the range of scores occupied by TD individuals^23^; both suggest that behind case-control comparisons may be additional heterogeneity nested within the ASC population. However, a problem in determining whether subgroups can be delineated on a measure like the RMET is that the canonical output of the test is one-dimensional, and would require experimenter-derived cut-points that would ultimately be arbitrary and not well informed about whether such cut-points are natural dividing points for stratification that reflect quantitatively and/or qualitatively different subgroups.

Here we overcome this problem by employing a novel data-driven subgrouping approach on item-level performance patterns on the RMET in two relatively large samples. Since we have little prior knowledge regarding how subgroups might manifest as item-level patterns of performance, we have opted for a data-driven unsupervised approach to stratification to open up new knowledge about how any such subgroups are present within performance on the RMET. Our stratification approach is an unsupervised hierarchical clustering procedure traditionally applied to high-throughput data generated in the fields of genomics and systems biology in similar situations where the experimenter typically has impoverished knowledge or little motivation to exert preconceptions or assumptions about precise distinctions or organization present in the data and whereby the goal is to have the data itself naturally present its organization. We show the existence of 5 separate ASC subgroups and 4 separate typically-developing (TD) subgroups that replicably appear across two large independent datasets. ASC subgroups separate into divisions that show very large differences compared to TD subgroups, as well as other ASC subgroups that show little to no difference. Thus, our unbiased data-driven stratification approach enhances the precision with which we can make more individualized statements about subsets of the ASC population while at the same time provides a novel methodological approach that is free from arbitrary experimenter-derived criteria and other potential biases that may affected supervised approaches and which may uncover new aspects of organization within the ASC population that have not been previously considered.

## Results

Our clustering approach discovered 5 distinct ASC subgroups (Fig 1A–B) and 4 distinct TD subgroups (Fig 1C–D) that replicably appear in both the Discovery and Replication datasets. Upon computing RMET sum scores for each subgroup, it is clear that clustering identifies natural subgroup divisions in the data that reflect different patterning of responses that result in quantitative differentiation in overall performance. Rank ordering the subgroups by RMET total scores results in a near linear trend for increasing RMET performance (Fig 2A–B). In characterizing effect sizes for ASC versus TD subgroup comparisons (Fig 2C–D) it is clear that the subgrouping procedure achieves the goals of enhancing sensitivity for identifying discrete ASC subgroups with large deficits, while at the same time enhancing specificity by showing that there are other ASC subgroups that show no sign of difficulty and are within the range of scores observed in the TD population. For example, the two poorest-performing ASC subgroups (subgroups 1 and 2) were dramatically lower than the range of scores observed in any of the TD subgroups with an effect size reduction in performance greater than 1.20 standard deviations. In the most extreme case when the worst performing ASC subgroup (ASC subgroup 1) was compared to the best performing TD subgroup (TD subgroup 4), the effect size rose to as high as 11.21 and 8.97 standard deviations respectively for each dataset. These poorest-performing ASC subgroups composed a relative minority of the Discovery (19%) and Replication (36%) ASC samples. ASC subgroup 3 is an intermediate subgroup, whereby inferences depend on which TD subgroup they are compared to. This subgroup is within the lower range of scores typically seen in the TD samples, and thus would show no deficit compared to the poorest-performing TD subgroup (Replication Cohen’s d = 0.15) or even slightly enhanced performance (Discovery Cohen’s d = 0.79). The poorest-performing TD subgroup was however, a minority of the TD samples comprising only 19-22% of individuals. Thus, in comparison to any of the other TD subgroups representing the majority of TD individuals (77-80%), the intermediate ASC subgroup 3 still shows pronounced deficits greater than 1 standard deviation of difference. Combining the clearly impaired (subgroups 1-2) and intermediate (subgroup 3) ASC subgroups captures 45-62% of the ASC individuals in each dataset respectively. ASC subgroups 4 and 5 comprise the remaining ASC individuals, who showed performance well within the range of scores observed in TD subgroups. However, even amongst these higher performing ASC subgroups, there was evidence for on-average differentiation when compared to the highest performing TD subgroups. For example, the effect size for a relatively highest-performing ASC subgroup (subgroup 4) versus the highest-performing TD subgroup (subgroup 4) was still a reduction around 2.13 and 2.58 standard deviations respectively. In contrast, there was also evidence that some high-performing ASC subgroups performed much better than the poorest-performing TD subgroups (e.g., ASC subgroup 5 vs TD subgroup 1 Cohen’s *d* = 3.72 and 2.33) (Fig 2C–D). This evidence of complex nested heterogeneity both within the ASC and TD samples points towards the need to make subgroup distinctions. Inferences without such important distinctions (i.e. standard case-control comparisons) could be biased in either direction depending on the mixture of heterogeneous individuals from various ASC and TD subgroups that would be nested in any one study.

**Figure 1:**
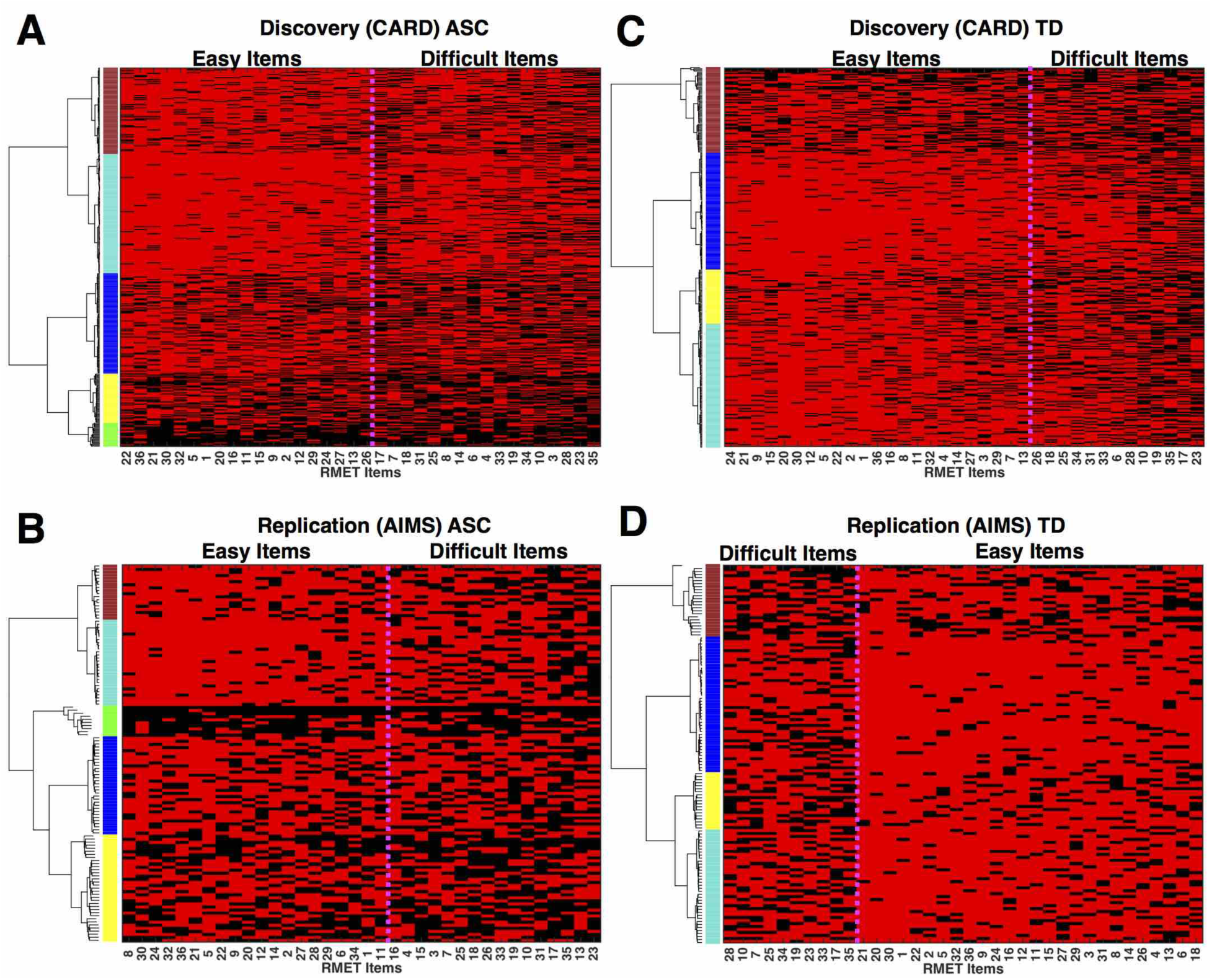
Clustering individuals into subgroups by RMET performance. This figure illustrates the raw data from both Discovery (A, ASC; C, TD) and Replication (B, ASC; D, TD) datasets in a 2D matrix (i.e. rows are subjects, columns are RMET items). Red cells indicate correct responses. Black cells indicate incorrect responses. The rows (subjects) and columns (items) of each matrix are rearranged by hierarchical clustering on topological overlap similarity in RMET response patterns. Subgroups are denoted by the different colors underneath the dendrogram branches.

**Figure 2:**
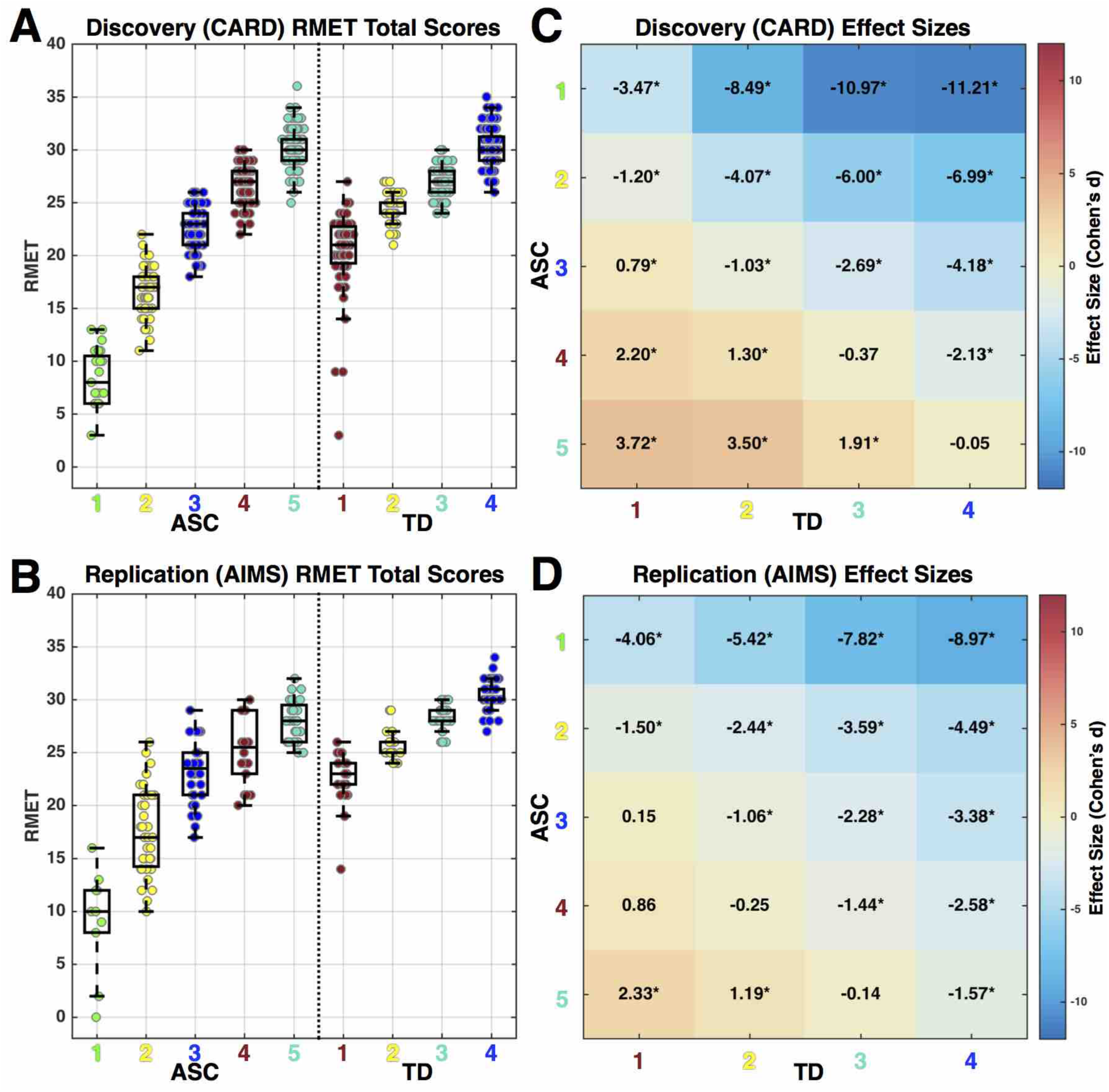
RMET total scores and effect sizes for comparisons across ASC and TD subgroups in Discovery and Replication datasets. Panels A and B show RMET total scores for each ASC and TD subgroup as boxplots along with dots overlaid to represent individual subject’s data points (Panel A depicts the Discovery (CARD) dataset, panel B depicts the Replication (AIMS) dataset). Panels C and D show standardized effect sizes (Cohen’s d; mean difference in units of standard deviation) for all pairwise comparisons of ASC versus TD subgroups. The effect sizes can be seen numerically within each cell and are also depicted by the coloring of the cell. The directionality of effect sizes can be interpreted as follows: negative values indicate an effect of TD subgroup > ASC subgroup, positive values indicate an effect of ASC subgroup > TD subgroup. An asterisk represents comparisons that pass Bonferroni correction for 20 pairwise comparisons.

Although ASC subgroups show item-level patterning that is reflective of quantitative differentiation in overall performance, it is not the case that signal reflected in quantitative differences in overall performance drives all of the differentiation between subgroups. Rather, subgroups also show dissimilar patterning of item difficulty across items. To better understand variability at the level of items, we ran identical clustering procedures (i.e. Ward hierarchical clustering on topological overlap) along the item dimension. The top two cluster branches can be described as differentiation between easy versus difficult items (Fig 1 and Fig 3). To test for similar or different patterns of item difficulty across the subgroups, we first computed a measure of item difficulty, defined as the percentage of individuals in a particular subgroup who answered a specific item correctly. Item difficulty plots for each ASC subgroups can be seen in Fig 3A–B (for similar plots on the TD subgroups see Supplementary Figure 1). Next, we assessed similarity in item difficulty by computing correlations across all pairwise subgroup comparisons, separately for subsets of easy or difficult items. Higher correlations indicate that item difficulty patterns are similar between subgroups, whereas lower correlations indicate more dissimilarity of item difficulty across subgroups. Here we find some evidence for similar patterning of item difficulty between ASC subgroups, restricted primarily to comparisons of certain adjacent rank-ordered subgroups, particularly subgroups 3-5. However, these correlations did not easily translate to replicable effects across both the Discovery and Replication datasets, and in the case of difficult items in the Replication dataset, no significant item difficulty correlations emerged. The general lack of replicable significant correlations between subgroups in item difficulty indicates that most pairwise subgroup comparisons are not highly similar in item difficulty patterns. This result suggests that in addition to picking up subgroups that can be characterized by quantitative differences in overall performance, the clustering approach also leverages useful information in the patterning of performance at the item-level to identify discrete subgroups.

**Figure 3:**
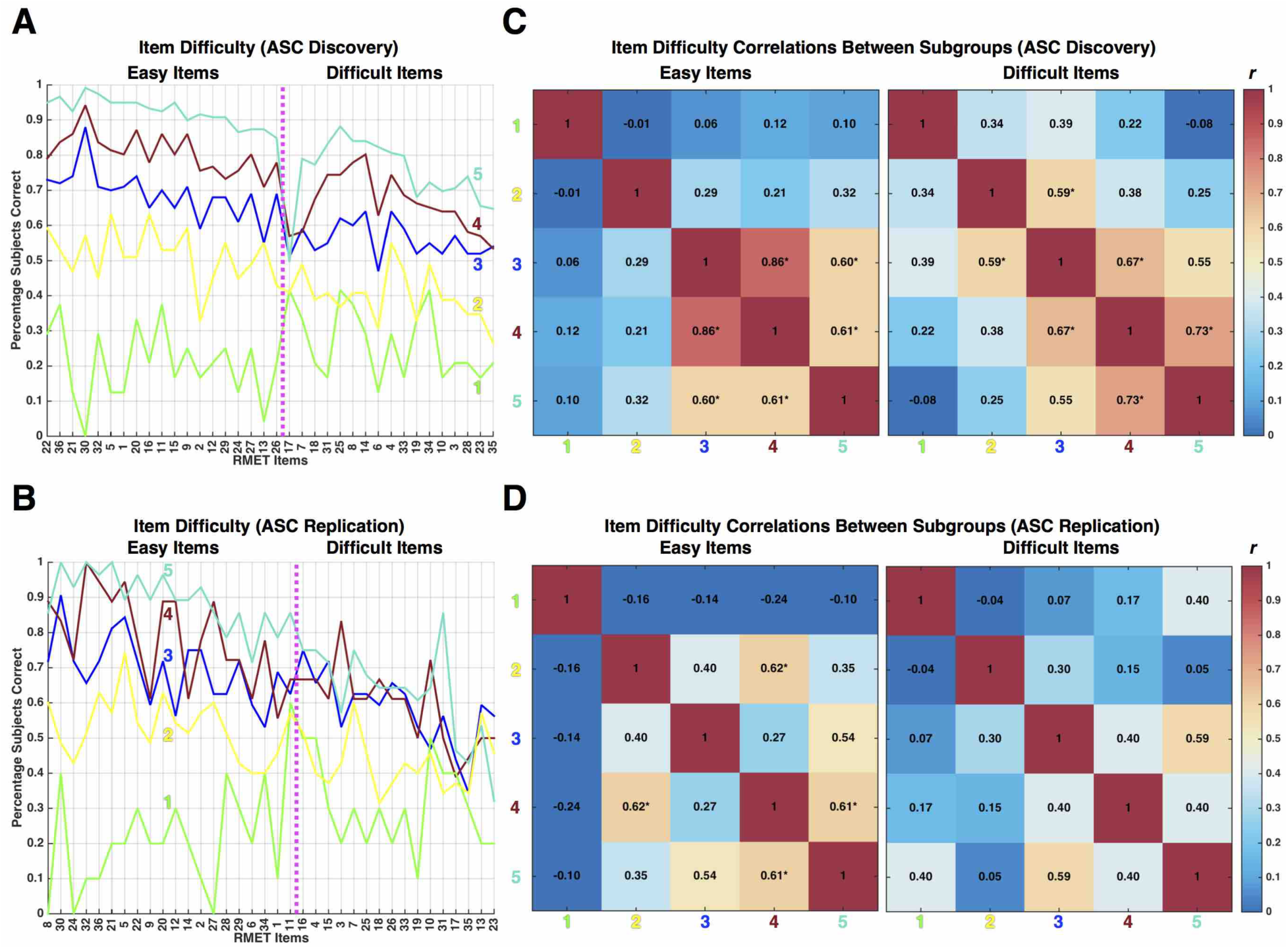
Item difficulty patterning across ASC subgroups. Panels A (ASC Discovery) and B (ASC Replication) show item difficulty profiles (i.e., percentage of subjects within a subgroup that answer the item correctly) for each ASC subgroup denoted by the different colored lines. Panels C and D show correlation matrices from item difficulty between subgroups. Asterisks indicate specific comparisons that pass FDR q<0.05 correction for multiple comparisons.

Next, we asked the question of whether individuals within subgroups are highly homogeneous in item-level patterning and whether similarity extends across the independent datasets. To begin exploring this issue, the Discovery and Replication subgroups were concatenated in rank-order along the subject dimension into one large two-dimensional matrix (subjects x items) and we then computed the full distance matrix across all subjects. This procedure was done for subsets of easy and difficult items separately. These subject-wise distance matrices allow for visualization of the full spectrum of between-subject dissimilarities within and across subgroups and across different datasets (Fig 4A–B; for TD matrices see Supplementary Figure 2). One overall gradient pattern emerges immediately in visual comparison across both easy and difficult item subsets. There is a general trend for marked between-subject similarity within subgroups at the poles of our rank-ordered subgroup hierarchy. That is, the worst and best performing subgroups tend to show high degrees of similarity within the subgroup boundaries (denoted by dark blue coloring in Fig 4) and this effect generalizes to the homologous rank-ordered subgroup in the other independent dataset. While this effect tends to be most pronounced for the easy item subset, it can also be seen in the difficult item subset. Between-subject similarity decreases in a gradient fashion as one descends down the rank-ordered subgroup hierarchy from ASC subgroup 5. ASC subgroup 2 shows the highest levels of between-subject dissimilarity that hover around mid-range Hamming distance values of 0.5 to 0.6 that denote 50-60% of items are dissimilar responses. This indicates that item-level patterning for individuals within ASC subgroup 2 are slightly dissimilar, relative to the much higher degree of similarity observed in other subgroups. This effect may be reflective of more random patterns of correct and incorrect responses, rather than consistent patterns of responses that are more homogeneous across all individuals within that subgroup.

**Figure 4:**
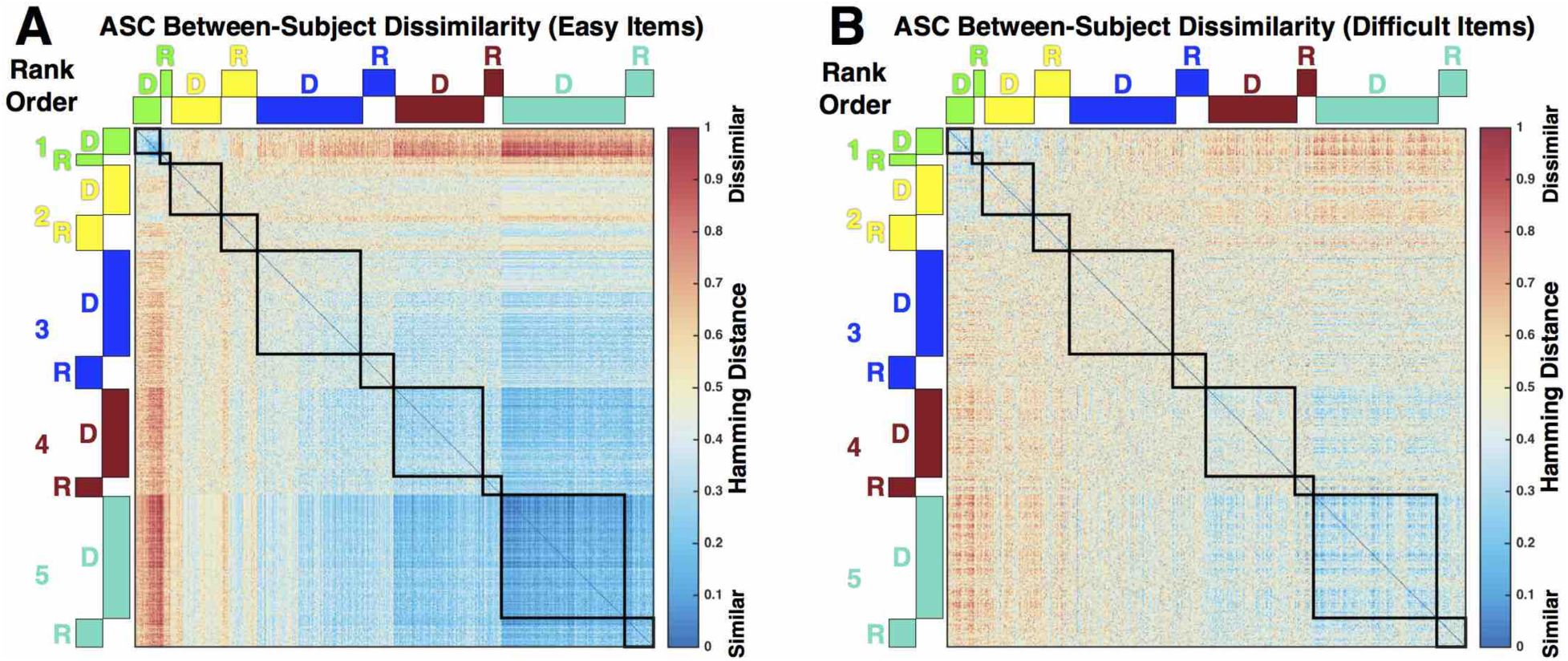
ASC Between-subject dissimilarity of RMET response patterns. This figure depicts between-subject dissimilarity matrices in ASC for the easy item (A) or difficult item (B) subsets. Cooler colors indicate more between-subject similarity, whereas hotter colors indicate more between-subject dissimilarity. Each cell of the matrices represents the dissimilarity between a pair of subjects. The rows and columns are arranged by subgroup rank order and Discovery and Replication datasets are adjacent to each other and denoted above the rows and columns by D and R. The black outlines delineate between-subject dissimilarities within a particular subgroup.

To go further in specifically testing whether subgroup divisions in one dataset generalize to independent data, we used multi-class classification analyses. This analysis allows for an explicit test of the hypothesis that the subgroups identified in one particular dataset are not simply reflecting idiosyncratic variability for that specific dataset but rather indicate generalizable subdivisions within the population structure of ASC. A 5-way classifier on ASC subgroups achieves 65% accuracy, which is never otherwise observed within 10,000 permutations of subgroup labels (p<9.99e-5; range of classification accuracy in the null distribution: 12.69% to 33.70%) (for TD classification see Supplementary Figure 3; for classification null distributions see Supplementary Figure 4).

We then inspected the confusion matrices for such multi-class predictions and this illuminated further key considerations. The misclassifications were nearly always for proximal (i.e. adjacent in rank-order) subgroups. An important distinction could be made between subgroups 1 and 2 versus subgroups 3-5. Nearly all the predictions for individuals within subgroups 1-2 or 3-5 stay within such superordinate groupings. Given that there was an important distinction between subgroups 1-2 being the most profoundly affected or ‘impaired’ ASC subgroups, whereas subgroups 3-5 showed performance within the TD range, it is interesting to consider that the accuracy recalculated for a distinction between subgroups 1-2 versus subgroups 3-5 is 92% and 93% respectively (Fig 5). Expanding the set of ‘impaired’ subgroups to also include the intermediate subgroup 3 and comparing them to subgroups 4-5 results in 88% and 84% accuracy respectively. This indicates that while the much harder task of multi-class prediction is not perfect, a coarser stratification into what could be considered roughly as ‘impaired’ versus ‘intact’ subgroups is much more robust.

**Figure 5:**
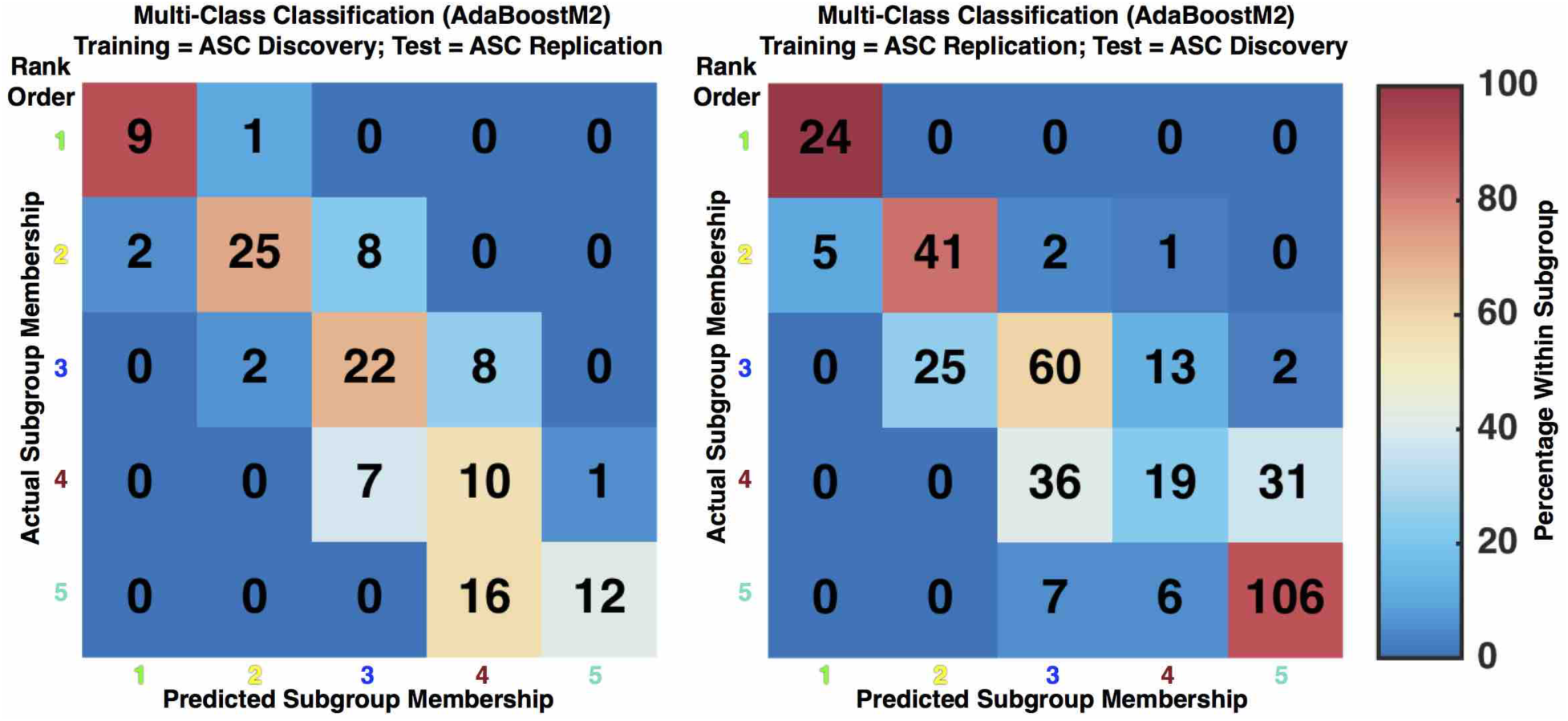
Confusion matrices for multi-class classifier predictions of ASC subgroup membership. Confusion matrices show the counts of actual ASC subgroup membership along the rows and classifier predicted subgroup membership along the columns. The coloring of cells in the confusion matrices represents the percentage of actual subgroup individuals predicted within each subgroup category. Above the matrices are descriptions of which dataset was used for training and testing.

Finally, we examined whether ASC subgroups differed on other variables such as sex/gender, age, AQ, EQ, BDI, BAI, ADOS and ADI-R scores. ASC subgroups did not systematically differ across both Discovery and Replication datasets by sex/gender or EQ (see Supplementary Table and Supplementary Figures 5-9). Although there were significant age effects in both datasets (Discovery: *F(4,373)* = 2.55, *p* = 0.03; Replication: *F(4,118)* = 3.48, *p* = 0.009), these effects were driven by different pairwise subgroup comparisons and were thus not systematic across the datasets. Within the AIMS dataset, where depression (BDI), anxiety (BAI), and autism symptom severity data (ADOS, ADI-R) were also available, no differences emerged between the subgroups (see Supplementary Table and Figures). Subgroups did show some important differences on the AQ and VIQ. For AQ, ASC subgroup differences manifested in both Discovery (*χ*^2^*(4,373)* = 13.69, *p* = 0.008) and Replication (*χ*^2^*(4,104)* = 9.89, *p* = 0.04) cohorts with the most RMET-impaired ASC subgroup (i.e., subgroup 1) showing markedly higher AQ scores than other better performing ASC subgroups (Fig 6A–D). For VIQ there was a clear effect of lower VIQ in poor RMET performing ASC subgroups(*F(4,118)* = 3.76, *p* = 0.006; Fig 6E–F). Given the presence of such an effect, we re-ran all hypothesis tests on RMET between-group differences while controlling for variability in VIQ and came to identical conclusions regarding robust differences. Therefore, although VIQ was lower in poor performing ASC subgroups, this could not be considered the primary explanation behind the subgroups’ poor RMET performance.

**Figure 6:**
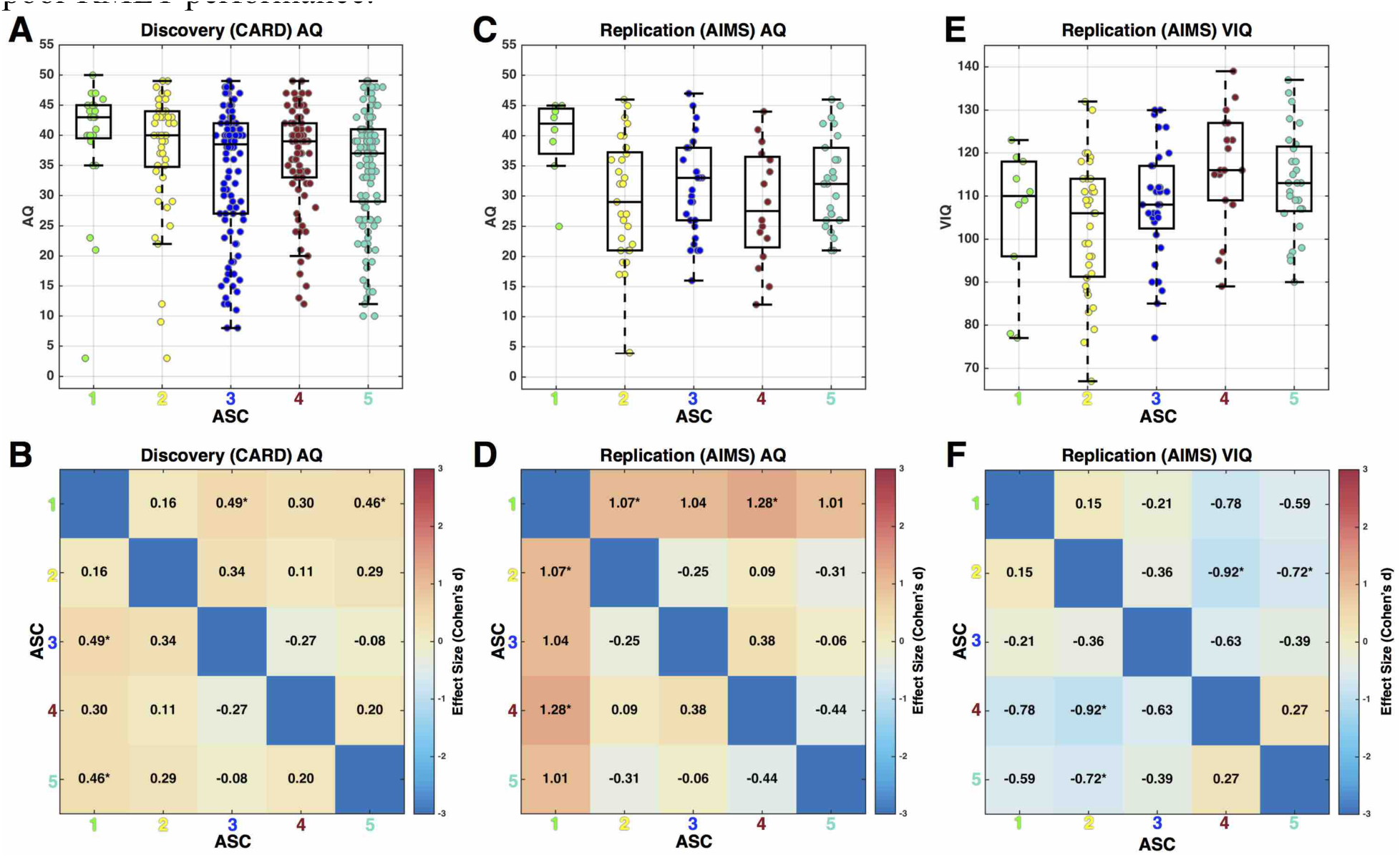
Characterization of subgroup differences on autistic traits (AQ) and verbal IQ (VIQ). Panels A, C, and E are boxplots with dots overlaid to show individual subject’s data points. Panels B, D, and F are heatmaps depicting the effect size for ASC subgroup comparisons. AQ data within the Discovery (CARD) dataset are shown in panels A and B, whereas AQ data within the Replication (AIMS) dataset are shown in panels C and D. Panels E and F show VIQ data from the Replication (AIMS) dataset. Effect sizes are standardized effect sizes (Cohen’s d) and interpreted as the mean difference in units of standard deviation.

## Discussion

In this study we examined heterogeneity in mentalizing ability as measured by the RMET in adults with and without ASC. We discovered 5 ASC subgroups and 4 TD subgroups that consistently emerged across two relatively large independent datasets. Variability in ASC ranged from subgroups that were very impaired to those within the TD range of scores, and in some cases, high-performing ASC subgroups were better than poor-performing TD subgroups. The spectrum of effect size differences between ASC and TD subgroups was highly consistent across independent datasets. The fact that replicable patterns of heterogeneity were found across datasets and across both TD and ASC underscores the idea that parsing heterogeneity amongst both ASC and TD populations is of considerable importance. The enhancement of sensitivity and specificity over and above a standard case-control comparison is self-evident when comparing the effect sizes in Figure 2C–D to the case-control effect sizes of Cohen’s d = −0.36 and −1.15 across Discovery and Replication datasets respectively. Such case-control comparison effect sizes mask a large degree of complexity hidden within both ASC and TD populations. Therefore, the precision at which we can understand mentalizing in ASC may be limited until we gain a better grasp on the nested heterogeneity present in such a domain both within the ASC and TD populations.

It is particularly noteworthy that the two poorest performing ASC subgroups (i.e. subgroups 1 and 2) were a relative minority of the sample in both datasets (i.e. 19% of the Discovery (CARD) dataset and 36% of the Replication (AIMS) dataset). When these ASC subgroups are considered relative to the other subgroups within the TD-range of scores (i.e. ASC subgroups 3-5), ability to accurately make a binary ‘impaired’ versus ‘intact’ prediction from a multi-class classifier is very high (92-93%). ASC subgroup 3 also represents an intermediate degree of impairment, as this subgroup shows decreased scores compared to all but the poorest-performing TD subgroup comprising the bottom 19-22% of TD individuals. Including ASC subgroups 1-3, the estimates of ASC individuals showing subtle-to-large impairment on the RMET ranges from 45-62%. Therefore, unlike the case of earlier points in development where a large percentage of individuals show difficulty in the domain of mentalizing^21^, in adulthood many individuals do not fall into subgroups 1-3 that would be considered impaired. One important caveat in interpretation is that these numbers are based on one test of mentalizing (i.e., the RMET). Clearly, there are other ways to measure different components of mentalizing ability and the RMET likely only taps specific components within the larger umbrella of mentalizing^22^. Examination of mentalizing subgroups using other tests will be particularly helpful for better understanding how heterogeneity manifests in similar or different ways across different aspects of mentalizing. Another caveat is that the RMET may not be sensitive enough to detect more subtle difficulties that translate better into understanding of real-world naturalistic social interactions, or which may be better measured with tasks/tests that tap an individual’s ability to automatically or implicitly mentalize, or which employ utilization of mentalizing to make more complex judgments (e.g., moral judgments)^19, 20, 25, 26, 27, 28, 29, 30^. Nevertheless, our findings of discrete, replicable, and robust ASC subgroups with differing explicit mentalizing ability as measured by the RMET in adulthood represents an important stride forward in terms of the precision of our understanding of mentalizing difficulties in adults with ASC. This work is highly compatible with the goals of ‘precision medicine’ or ‘stratified psychiatry’ and is what is needed to move forward with research that has clinical impact for patients and which can also further translational research progress focused on honing in on treatment-relevant mechanisms^12, 14^

The subgroup distinctions outlined here are also particularly important because of how they apply specifically to the RMET. The RMET is a long-standing instrument that is widely used within the fields of autism research and social neuroscience. The NIMH Research Domain Criteria (RDoC) lists the RMET as one of several important tests for characterizing variation in social processes, particularly under the category of Perception and Understanding of Others (http://Lusa.gov/1Qs6MdI). In addition to its wide usage in autism research, the RMET has also been used to characterize and compare social-cognitive abilities across different categorical psychiatric diagnoses ^31, 32, 33, 34^. With regards to treatment research, the RMET is widely used as treatment outcome measure, particularly for drug manipulations (e.g., oxytocin) or behavioral interventions targeting social skills and social cognition ^35, 36, 37, 38^. All of this prior clinically important research utilizes an analytic strategy of computing RMET summary scores across all items and then onto potentially sub-optimal omnibus case-control comparisons that may mask the presence of nested subgroups within ASC. The current work should signal a change in this practice for how the RMET is utilized in important clinical settings (e.g., evaluating treatment outcome). Rather than using summary scores in an omnibus ASC group, a more fruitful approach would be to use the RMET to distinguish subgroups and to then specifically evaluate whether such ASC subgroups respond differently to treatment. In other words, the added knowledge we provide here is that these subgroups could signal a meaningful distinction that helps in the design of intervention studies and the subsequent interpretation of such findings. Given the current state of largely mixed results for many interventions for ASC^39^, it may become clearer after subgrouping that some treatments do systematically work for particular subgroups but not others.

In addition to impact in clinical research areas, the current study could potentially have large impact on basic research targeting mechanisms and the phenotypic diversity of ASC. For example, inconsistency within the literature on the cognitive phenotype of ASC^40^, particularly as it pertains directly to mentalizing and/or more generally the domains of emotion and social cognition, may be better understood with an approach that focuses more on parsing heterogeneity into subgroups as we have shown here. Additionally, inconsistency in the functional and structural neuroimaging literature on ASC^41, 42, 43, 44, 45^ could be mitigated by a better understanding of mentalizing heterogeneity nested within relatively small ASC samples typically utilized in such work. This point is underscored by recent work by Byrge and colleagues, whereby it was suggested that some group-level differences in case-control designs could be driven by the effects nested within a small subgroup of patients^30^. A better a priori understanding of the heterogeneity present within the ASC population could be of large impact for study design and could also implicate different underlying etiological, neurobiological, and developmental mechanisms that explain such heterogeneity^7, 46, 47, 48^.

A major innovation in this work is the approach to subgrouping. Rather than utilizing the RMET in a standard approach by summarizing all items into one total score, we have instead retained the full set of information encoded across the 36 items as input into an *unsupervised* hierarchical clustering approach that came to data-driven conclusions about the presence of discrete ASC subgroups. This unsupervised approach avoids using potentially arbitrary experimenter-derived cutoffs and instead utilizes natural data-driven distinctions that are robust enough to emerge in a consistent fashion across independent datasets. Importantly, this clustering approach leverages distinctions in the patterning of item-level performance that can relate to quantitative distinctions in overall performance but also subtle dissimilarity in item difficulty across the subgroups. Recent work has applied similar logic and approaches to clustering the phenotype based on gold-standard diagnostic instruments^49^ and for clustering 26 different mouse models of genetic mechanisms related to ASC^46^. However, these other types of clustering approaches do not benefit from some of the specific innovations inherent in the technique we have used. Specifically, we utilized important computational steps taken directly from weighted gene co-expression network analysis (WGCNA)^50, 51^, which is a widely used approach in genomics and systems biology and has been highly utilized specifically in autism genomics research^52, 53, 54, 55, 56, 57^ In particular, the computational steps of converting a distance matrix into a topological overlap matrix and then running hierarchical clustering on this similarity metric rather than other metrics is important since topological overlap is less susceptible to noise influences because it leverages information about similarity of neighbors. Additionally, the dynamic hybrid tree-cutting algorithm that cuts the cluster tree into discrete subgroups is also highly innovative compared to most other tree-cutting methods which rely on using a single cut height across the dendrogram, and which generally cannot make fine distinctions that the dynamic algorithm can make within local neighborhoods of the dendrogram. Thus, our analytic approach is applicable across translational research contexts and could be utilized more widely across a whole range of new applications focused on data-driven stratification in ASC.

A further advantage to our approach of finding data-driven distinctions is that such distinctions are generalizable across datasets. As we have shown with the classification analyses, the stratifications made in one dataset generalize to multi-class predictions in independent data. In the context of a more simplistic 2-class distinction of ‘impaired’ versus ‘intact’, the multiclass predictions were even more accurate. In future work, such information about replicable subgroups could be turned into valuable assessment or research tools that could aid in study design and participant screening. For instance, randomized control trials may use the RMET to screen patients along such distinctions or as an outcome measure and use such distinctions to analyze individualized treatment response patterns. Such stratifications could also be useful in clinical assessments and facilitate personalized treatment planning and outcome prediction.

In addition to highlighting the promise of such stratifications for autism research, there are additional characteristics of ASC subgroups that are important to stress. First, variables such as sex/gender, age, trait empathy, depression and anxiety symptoms, and clinical measures of autistic symptom severity were not systematically different across ASC subgroups. However, poor performing ASC subgroups tended to be lower in VIQ and higher in self-reported autistic traits measured by the AQ. This effect of higher self-reported autistic traits in more affected subgroups is interesting from the standpoint that similar effects do not emerge on clinical measures of autistic symptom severity (e.g., ADOS and ADI-R). It may be that this effect emerges due to differences in what is measured in an instrument like the AQ versus the ADOS and ADI-R. This effect may be of clinical importance as those individuals with high levels of autistic traits and poor performance on the RMET may potentially need different approaches of intervention and support than other individuals within other subgroups (e.g., they may particularly benefit from adjustments in the occupational or educational environments to reduce social load). The VIQ effect on the RMET and more generally on mentalizing ability has been noted before ^21, 58^ and may be easily understood in the context of the RMET since this test may tax vocabulary for some individuals and on certain items. Despite this effect of VIQ, we found that the main comparisons of ASC versus TD subgroups were unchanged after accounting for VIQ variability. This evidence suggests that while some variability in RMET performance is linked to variability in VIQ, the subgroup distinctions and patterns of RMET performance are not fully explained by VIQ variation.

In conclusion, the discoveries in this study allow for a more precise understanding of mentalizing in adults with ASC. Our insights have the potential to further personalized medicine aims in ways that accelerate progress towards clinical impact for patients. By understanding how the autism spectrum can be stratified in clinically meaningful ways, translational opportunities may open up that could test whether such distinctions are rooted in separate underlying mechanisms.

## Materials and Methods

### Discovery Dataset

In this study we analyzed two large datasets that served as discovery and replication sets. The discovery dataset came from the Cambridge Autism Research Database (CARD)^23^ and consisted of 395 adults with ASC (178 males, 217 females) and 320 typically-developing controls (TD; 152 males, 168 females) within the age range of 18-74 years. The CARD data were collected online from two websites (www.autismresearchcentre.com,www.cambridgepsychology.com) during the period of 2007-2014. Once participants had logged onto either site, they consented for their data to be held in the Cambridge Autism Research Database (CARD) for research use, with ethical approval from the University of Cambridge Psychology Research Ethics Committee (reference No. Pre.2013.06).

CARD participants who self-reported a clinical autism diagnosis were asked specific information about the date of their diagnosis, where they were diagnosed, and the profession of the person who diagnosed them. The inclusion criterion for participants in the ASC group was a clinical diagnosis of an autism spectrum condition (ASC) according to DSM-IV (any pervasive developmental disorder), DSM-5 (autism spectrum disorder), or ICD-10 (any pervasive developmental disorder) from a recognized specialist clinic by a psychiatrist or clinical psychologist. Such online self or parent-reported diagnoses agree well with clinical diagnoses in medical records ^59^. Control group participants were included if they had no diagnoses of ASC and no first-degree relatives with ASC. For both groups, participants were excluded if they reported a diagnosis of bipolar disorder, schizophrenia, eating disorder, obsessive-compulsive disorder, personality disorder, epilepsy, or an intersex/transsexual condition. Participants with a diagnosis of depressive or anxiety disorder were not excluded as these conditions are common in the general population and occur at high rates in adults with autism^1^.

### Replication Dataset

The replication dataset consisted of participants from the MRC AIMS Consortium dataset (n=123 ASC; 85 male, 38 female; n=128 TD; 87 male, 41 female) within the age range of 18-52^60, 61, 62, 63^. The study was given ethical approval by the National Research Ethics Committee, Suffolk, UK. All volunteers gave written informed consent. Participants were recruited and assessed at one of the three MRC AIMS centers: the Institute of Psychiatry, London; the Autism Research Centre, University of Cambridge; the Autism Research Group, University of Oxford. All participants were right-handed. Exclusion criteria for all participants included a history of major psychiatric disorder (with the exception of depressive or anxiety disorders), head injury, genetic disorder associated with autism (e.g., fragile X syndrome, tuberous sclerosis), or any other medical condition affecting brain function (e.g., epilepsy). All ASC participants were diagnosed according to ICD-10 research criteria for pervasive developmental disorder. ASC diagnoses were confirmed using the Autism Diagnostic Interview-Revised (ADI-R)^64^ and it was allowed for participants to be 1 point below cutoff for one of the three ADI-R domains in the diagnostic algorithm. The Autism Diagnostic Observation Schedule (ADOS)^65^ was used to assess current symptoms for all participants with ASC. The Wechsler Abbreviated Scale of Intelligence (WASI)^66^ was used to assess Verbal IQ (VIQ), Performance IQ (PIQ) and Full Scale IQ (FSIQ). Depressive and anxiety symptoms were measured with the Beck Depression Inventory (BDI) and Beck Anxiety Inventory (BAI).

### Reading the Mind in the Eyes Test (RMET)

All participants in both discovery and replication datasets completed the ‘Reading the Mind in the Eyes’ Test (RMET), adult version^24^. The RMET consists of 36 items of grey-scale photos cropped and rescaled so that only the area around the eyes can be seen. Each photo is surrounded by four mental state terms and the participant is instructed to choose the word that best describes what the person in the photo is thinking or feeling. Participants in both discovery and replication datasets completed a computerized online version of the RMET at home. Participants were instructed to select the most appropriate item within 20 seconds for each stimulus (presented in random order). Responses were coded as correct or incorrect (wrong items selected, or no response after 20 seconds), giving a maximum total correct score of 36. To guard against the possibility that many items timed-out, we used a rule that if an individual had timeouts on 9 or more items (>25% of all items), then such individuals were excluded from analysis. All participants in both discovery and replication datasets also completed the Autism Spectrum Quotient (AQ)^67^ and the Empathy Quotient (EQ)^68^ on the same online platform and before taking the RMET.

### Statistical Analysis

RMET item-level data for all subjects was concatenated into a two-dimensional matrix with subjects along the rows and RMET items along the columns. This data matrix was then converted into a distance matrix across subjects. The value within each cell of this distance matrix indicates how similar each individual is to another individual in RMET item-level patterns of response. The distance metric computed was Hamming distance, which is a measure of the percentage of dissimilar item responses between two subjects and is appropriate in this context where RMET item-level responses are binary. For the purpose of clustering into subgroups, the distance matrices for each dataset were converted into a topological overlap matrix (TO). Topological overlap is an advantageous metric of similarity over and above other distance metrics that only take into account similarity between the two individuals of interest because topological overlap will also take into account similarity between the neighbors of the target individuals. When two individuals are highly similar between themselves and also in their neighbors, they have high topological overlap. Topological overlap matrices are highly effective in other applications^69, 70^ including the systems biology method of weighted gene co-expression network analysis (WGCNA)^71^. The topological overlap matrices were then input into agglomerative hierarchical clustering using Ward’s method as the linkage method. The dendrograms created from clustering were then cut into subgroups using a dynamic hybrid tree cutting algorithm (deepSplit = 1) also commonly used in systems biology applications such as weighted gene co-expression network analysis^72^. This tree-cutting algorithm is optimal for finding subgroups as it finds a dynamic cut height for each branch of the dendrogram rather than using a single cut height for all branches. This entire subgrouping procedure was implemented on both the ASC and TD groups independently.

Once subgroups were defined, we computed total RMET scores (i.e. sum across all items) for each individual and ran independent samples t-tests to specifically compare the total score across all pairwise comparisons of ASC subgroups versus TD subgroups. Only comparisons that passed Bonferroni correction for 20 comparisons were considered significant. Standardized effect size for each comparison was also computed as Cohen’s *d*. All pairwise comparisons between ASC and TD are visualized as heatmaps showing standardized effect size (Cohen’s *d*) for each comparison. Note that we did not compute within-group comparisons because such comparisons would be circular given that the subgrouping (selection) and testing would be done on the same data.

In addition to stratifying the subject dimension we also applied clustering to the item dimension of the dataset. This clustering was done to primarily find the two major subdivisions of items that could be characterized as easy versus difficult items. These subsets of items were then used in further analyses that examined different patterning of item difficult across the subgroups. To measure item difficulty we calculated the percentage of individuals within a particular subgroup that answered a specific item correctly. To examine the hypothesis that subgroups show similar or different item difficulty profiles we computed correlations between the item difficulty measures for each pairwise subgroup comparison. Correlations were deemed significant if they passed an FDR q<0.05 threshold that corrects for multiple comparisons. These significant correlations indicate subgroup comparisons whereby item difficulty was significantly similar across the subgroups. The non-significant correlations are taken to be subgroup comparisons whereby there was no sufficient evidence to state that item difficulty profiles were similar across the subgroups.

To examine between-subject dissimilarity of item-level performance patterns between subgroups and across datasets, we computed subject-wise distance matrices. These matrices show the similarity metric of Hamming distance for each pairwise subject combination across both datasets. These matrices were computed separately for the easy and difficult item subsets. These matrices primarily serve a descriptive purpose to explicate all between-subject dissimilarities and to show how similar individuals from a particular rank-ordered subgroup are to the homologous subgroup identified in the other dataset. If homologous subgroups identified in different datasets are indeed highly similar, we expect to see high degree of between-subject similarity across datasets.

To quantitatively evaluate the degree to which subgroups identified within one dataset could be accurately predicted within a second independent dataset we ran multi-class classification analyses using the ensemble learning algorithm AdaBoostM2^73^ implemented within the fitensemble.m function in MATLAB R2015b (learner type set to ‘Discriminant’ and with 20 weak learners). Homologous subgroup labels were based on rank ordering of the subgroups by total RMET scores. These homologous subgroup labels allowed us to then evaluate how well multi-class classification performance was in identifying the same rank ordered subgroups across datasets. Classification accuracy was then compared to simulations where subgroup labels were randomly permuted 10,000 times, and p-values were computed as the percentage of times under randomly permuted labels that classification accuracy was as high or higher than accuracy obtained under the true subgroup labels. To visualize multi-class classification performance, we present confusion matrices illustrating the percentage of individuals within each subgroup that are predicted in each subgroup category. We also computed classification accuracy for specific subsets of subgroups combined that could generally be called ‘impaired’ versus ‘intact’; that is, subgroups 1-2 versus subgroups 3-5 and subgroups 1-3 versus subgroups 4-5.

Finally, we examined other variables such as sex/gender, VIQ, age, AQ, EQ, BDI, BAI, and ADOS and ADI-R subscales to test hypotheses about whether the ASC subgroups would differ on these variables. To test for the possibility of imbalances across the subgroups as a function of sex/gender, we counted up the number of males and females across all subgroups and compared them to expected counts derived from a chi-square test. To test VIQ, age, AQ, EQ, BDI, BAI, ADOS, and ADI-R score differences we used one-way ANOVAs to test for differences between ASC subgroups. Because AQ and EQ showed markedly skewed distributions, we ran a Kruskal-Wallis one-way nonparametric ANOVA instead of a parametric ANOVA. ANOVA results were followed up with post-hoc pair-wise comparisons that were Bonferroni corrected for multiple comparisons.

## Acknowledgments

This study was supported by the National Institute for Health Research (NIHR) Collaboration for Leadership in Applied Health Research and Care (CLAHRC) East of England at Cambridgeshire and Peterborough NHS Foundation Trust. The views expressed are those of the authors and not necessarily those of the NHS, the NIHR, or the Department of Health. This study was also conducted in association with the European Autism Interventions—A Multicentre Study for Developing New Medications (EU-AIMS) consortium; EU-AIMS receives support from the Innovative Medicines Initiative Joint Undertaking under grant agreement number 115300, resources of which are composed of financial contribution from the European Union’s Seventh Framework Programme (FP7/2007–2013), EFPIA companies, and Autism Speaks. This study was also supported by grants from the UK Medical Research Council (MRC) (G0600977), the Wellcome Trust (091774/Z/10/Z), and the Autism Research Trust (ART). M-CL and AR received support from the William Binks Autism Neuroscience Fellowship at the University of Cambridge. M-CL received support from the O’Brien Scholars Program within the Child and Youth Mental Health Collaborative at the Centre for Addiction and Mental Health and The Hospital for Sick Children, Toronto.

## Author Contributions

All authors, including the MRC AIMS Consortium, contributed to the design of the study, performed the research, and contributed to the collection of data. MVL analyzed the data. MVL, M-CL, BA, and SBC wrote the manuscript.

## Additional Information

ETB is employed half-time by the University of Cambridge and half-time by GlaxoSmithKline (GSK). None of the other authors have any conflicts of interest to declare.

## MRC AIMS Consortium Author List (Alphabetical Order)

Anthony J. Bailey^11^, Simon Baron-Cohen^3^, Patrick F. Bolton^10^, Edward T. Bullmore^8^, Sarah Carrington^11^, Marco Catani^10^, Bhismadev Chakrabarti^3^, Michael C. Craig^10^, Eileen M. Daly^10^, Sean C. L. Deoni^10^, Christine Ecker^10^, Francesca Happé^12^, Julian Henty^3^, Peter Jezzard^11^, Patrick Johnston^10^, Derek K. Jones^10^, Meng-Chuan Lai^3^, Michael V. Lombardo^3^, Anya Madden^10^, Diane Mullins^10^, Clodagh M. Murphy^10^, Declan G. M. Murphy^10^, Greg Pasco^3^, Amber N. V. Ruigrok^3^, Susan A. Sadek^3^, Debbie Spain^10^, Rose Stewart^11^, John Suckling^8^, Sally J. Wheelwright^3^, Steven C. Williams^10^, and C. Ellie Wilson^10^.

